# Addressing structural mentoring barriers in postdoctoral training: A qualitative study

**DOI:** 10.1101/2022.08.20.504665

**Authors:** W. Marcus Lambert, Nanda Nana, Suwaiba Afonja, Ahsan Saeed, Avelino C. Amado, Linnie M. Golightly

## Abstract

**Background:** Structural mentoring barriers are policies, practices, and cultural norms that collectively disadvantage marginalized groups and perpetuate disparities in mentoring. While these mentoring barriers can be found early in the training pathway, failure to address or overcome these barriers at the postdoctoral training stage has a direct impact on faculty diversity and national efforts to retain underrepresented groups in research careers.

**Methods:** To better understand the mentoring barriers faced by postdoctoral trainees, and possible ways to address them, a diverse sample of postdoctoral scholars (“postdocs”) from across the United States were asked to participate in focus groups to discuss their training experiences. We conducted five 90-minute focus groups with 32 biomedical postdocs, including 20 (63%) women and 15 (47%) individuals from underrepresented racial/ethnic groups (URG). Participants were well-represented across years of training, and 65% were at least somewhat likely to pursue a research-intensive faculty career, similar to previously reported national averages.

**Results:** A social ecological framework was used to examine both the upstream and downstream manifestations of structural mentoring barriers, as well as mentor barriers, overall. Themes were categorized on four broad levels: *Individual* (attitudes, beliefs, knowledge, or behaviors that inform mentoring barriers), *Interpersonal* (mentoring barriers arising from dyadic, peer, or network relationships), *Institutional* (departmental, institutional, organizational mentoring barriers), or *Systemic* (mentoring barriers originating from policies or broad social and cultural norms). Notable structural barriers included (1) academic politics and scientific hierarchy, (2) inequalities resulting from mentor prestige, (3) the (over) reliance on one mentor, (4) the lack of formal training for academic and non-academic careers, and (5) the lack of institutional diversity and institutional mentor training. These structural barriers foster mentoring practices and behaviors that lead to poor work-life balance, poor communication, and research career attrition. To overcome these barriers, postdocs strongly encouraged developing a network or team of mentors and recommended institutional interventions that create more comprehensive professional development, mentorship, and belonging.

**Conclusions:** For postdoctoral scientists, structural mentoring barriers can permeate down to institutional, interpersonal, and individual levels, impeding a successful transition to an independent research career. It has become clear that large-scale changes in mentoring must come from addressing the policies, practices, and cultural norms that perpetuate poor mentoring. This work provides strong evidence for promoting mentorship networks and cultivating a “mentoring milieu” that fosters a supportive community and a strong culture of mentorship at all levels.

## Introduction

Effective mentorship during the postdoctoral (“postdoc”) training stage is crucial for successfully transitioning to an independent research career. It has been shown that receiving effective mentorship has a significant positive effect on early-career researchers’ satisfaction, self-efficacy, and career outcomes [1–5]. Mentorship of trainees, specifically in areas of grant-writing, is positively associated with increased publication productivity and self-efficacy [6–9]. Mentoring in scientific communication skills increases research career intention [10, 11]. Furthermore, career coaching has been shown to effectively supplement traditional one-on-one research mentoring [12, 13].

Our previous work has shown that postdoctoral scholars from underrepresented groups (URGs) leave the academic research pathway in part due to perceptions of poor mentorship [14]. The likelihood of biomedical postdocs choosing an academic research career increases as financial security, mentorship from their PI, and their sense of self-worth increase. We have also codified advice from postdoctoral trainees on pursuing a research career in academia, and one of the top recommendations was finding a strong mentor [5]. Trainees from URGs do not often find mentors that match their identity, and it has been shown that higher levels of mentor-protégé psychological similarity are related to higher levels of psychosocial support and relationship satisfaction [15, 16]. Overall, mentorship at the postdoctoral stage remains key for increasing workforce and faculty diversity [17]. However, postdocs, especially from underrepresented groups (URGs), face several structural mentoring barriers that impact their career outcomes.

Structural mentoring barriers are policies, practices, and cultural norms that (intentionally or unintentionally) perpetuate inequities in mentoring for some groups based on their identity. Structural barriers often favor an advantaged group while systematically disadvantaging a marginalized group [18]. For example, structural barriers have led to racial inequities in R01 funding [19, 20]. However, structural *mentoring* barriers have exacerbated these outcomes. Unlike graduate students, postdoctoral researchers are not systematically assigned an advisory or mentorship committee. In addition, many institutions have not yet offered or required formal mentor training for faculty [21]. Some institutions do not have postdoc offices or administrative positions to support postdocs. This can collectively lead to postdocs who are underpaid and exploited by faculty research mentors [1, 5].

Moreover, faculty from URGs are often disproportionately tasked with service and mentorship responsibilities, known as a “minority tax” or “cultural taxation” [22–25]. Women and faculty from underrepresented racial/ethnic groups are asked to serve on institutional committees, engage in outreach and educational activities, diversity efforts, and mentor trainees much more so than their peers from well-represented groups (WRGs) [24, 25]. This mentorship imbalance can have significant effects on mentoring outcomes of trainees from URGs [26].

Structural mentoring barriers can also negatively impact career outcomes. Many PhD and postdoc trainees report very little guidance from research mentors for a broad range of career paths [14, 27]. This is often a result of structural barriers. Postdocs are exposed to minimal career development resources during their training, resulting in a lack of clarity, confidence, and skills to efficiently find the right career fit [28, 29]. Even when postdocs are offered structured career development resources (e.g., individual development plans (IDPs), and/or workshops on transferable skills training), the perceived usefulness of these sessions varies [30–33].

In this article, we report the results of focus groups that explore the mentoring barriers faced by postdoctoral trainees and possible solutions to dismantle those barriers. We hypothesize that there are a significant number of postdocs facing mentoring barriers that are structural in nature. Using a social ecological framework to examine mentoring barriers, we have learned: i) broad systemic mentoring barriers can be mitigated through policies, networks, and training; (ii) the lack of institutional diversity and mentor training are impacting postdoctoral mentoring relationships and career outcomes; and (iii) many postdocs, especially from URGs, seek more nurturing mentors who understand their values and can build their confidence.

## Methods

### U-MARC database

Postdoctoral scholars interviewed during this study were recruited from the U-MARC database, which was developed in 2017 from U.S. postdocs in the biological and biomedical sciences. These postdocs completed the U-MARC (Understanding Motivations for Academic Research Careers) survey instrument and volunteered their contact information for future studies. The 70-item survey measures factors that determine career choice in science [14]. Sampled postdocs in the database represent 6% of the total pool of appointed biomedical and biological postdocs (1,292 of 21,781) the year the survey was conducted (2017) according to the National Science Foundation (NSF). The sample represents wide, geographic (over 80 universities), and subfield diversity. The REDCap electronic data capture tool was used to collect and manage the database. REDCap (Research Electronic Data Capture) is a secure, web-based application designed to support data capture for research studies [34].

### Selection of study participants

Of the 1,292 original survey participants, 346 indicated that they were not interested in participating in a follow-up focus group at the time of the original survey (**Fig 1**). So, they were not included in this outreach. 946 postdocs were contacted and were invited to participate in a follow up study about academic research careers. Postdocs were asked to complete a pre-screener survey that captured emails, names, race, ethnicity, citizenship status, their current academic institution, number of years as a postdoc, and how likely they were to pursue a research-intensive faculty career. 130 respondents self-selected to join the focus group through the prescreening survey. All work was conducted under the approval of the Weill Cornell Medical College Institutional Review Board (IRB# 1612017849), and all candidates provided consent for participation in the study. All 130 respondents were invited to select time slots for the focus groups. We conducted five 90-minute focus groups with 32 postdocs, including 20 women (63%) and 15 individuals from underrepresented racial/ethnic groups (47%). Three focus groups were comprised of postdocs from underrepresented groups and two focus groups were comprised of postdocs from well-represented groups. All participants in the focus groups were awarded a $50 gift card as compensation for their time and voluntary participation.

**Fig 1.**
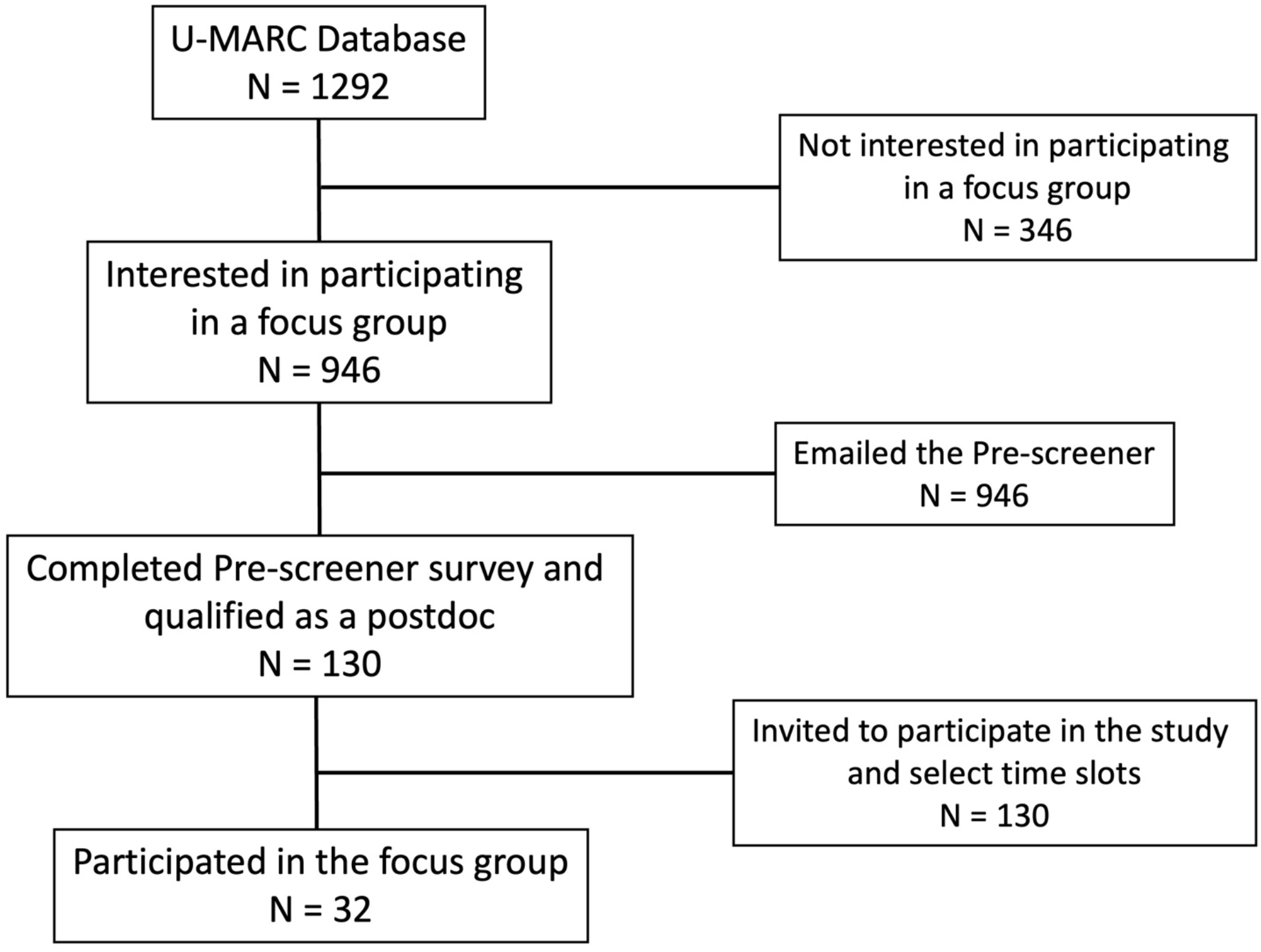
Flow Chart of Focus Group Participants. Flow diagram of the process for selection of focus group participants. Abbreviations: U-MARC, Understanding Motivations for Academic Research Careers.

### Interviews and Data analysis

Five focus groups were conducted virtually via Zoom with one moderator who guided the discussion and one co-moderator who observed facial expressions and helped to keep time. We began the focus group with introductions and a review of the study objectives and ground rules. We notified the participants to respect the privacy of the others and not repeat what was said in the focus groups. Participants were asked several open-ended questions from an interview guide that began with a discussion of their values and what they found most fulfilling in their work. We also asked, “what have mentors done to support or hinder your ability to pursue a research faculty career?” The study ended when data saturation was reached (the point at which no new information or themes were observed for the data). Each focus group was video recorded, after which the video was sent to an outside company for transcription. The transcripts of each session were stripped afterwards of any identifying information and each participant was assigned a participant ID number.

All the data were analyzed manually by an inductive method following these steps. First, each focus group was assigned to two researchers trained in qualitative coding. A process of open, axial, and then selective coding was followed by generally coding and discussing major concepts, categories, and themes. These themes and categories were inductively generated during the research, according to the grounded theory method. The two researchers involved in the coding process each independently derived codes, then met with a team of five to six researchers to help determine crosscutting themes and recurrent patterns. We repeated this cycle until we achieved thematic saturation, and novel themes stopped emerging from the data. NVivo 12, a qualitative transcript software, was used to assist with the coding of the data. Once all of the data were coded and themes emerged, we focused on the codes that dealt with aspects of mentorship, using an ecological systems approach to examine mentoring barriers.

## Results

To better understand the mentoring experiences of postdoctoral trainees, we interviewed a sample of 32 postdoctoral researchers across five 90-minute focus groups. 20 participants were women (63%), and 15 individuals were from NIH/NSF-designated underrepresented racial/ethnic groups (URG) (47%) (**Table 1**). These racial/ethnic groups include American Indian or Alaska Native, Black or African American, and/or Hispanic or Latino. Participants were well-represented across years of training, with 16% of participants in year 1 (less than 1 year of training), 22% in year 2, 25% in year 3, 16% in year 4, 13% in year 5, and 9% in year 6 or beyond. 65% of participants were at least somewhat likely to pursue a research-intensive faculty career, similar to previously reported national averages [14, 35]. Participants from URGs skewed towards pursuing a research-intensive faculty career (60% very likely).

**Table 1.**
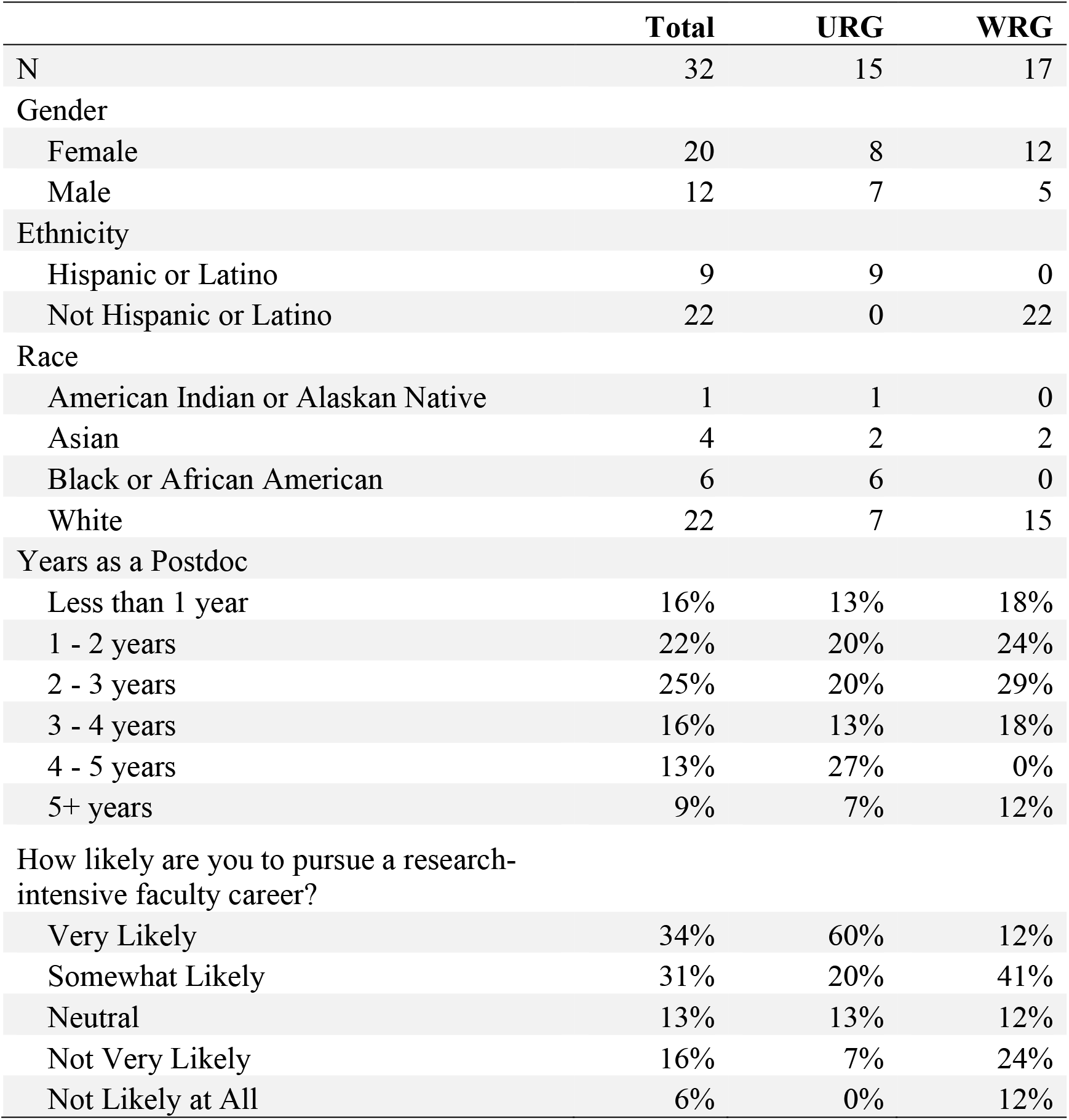
Demographics of Interview Participants.

|  | **Total** | **URG** | **WRG** |
| --- | --- | --- | --- |
| N | 32 | 15 | 17 |
| Gender |  |  |  |
| Female | 20 | 8 | 12 |
| Male | 12 | 7 | 5 |
| Ethnicity |  |  |  |
| Hispanic or Latino | 9 | 9 | 0 |
| Not Hispanic or Latino | 22 | 0 | 22 |
| Race |  |  |  |
| American Indian or Alaskan Native | 1 | 1 | 0 |
| Asian | 4 | 2 | 2 |
| Black or African American | 6 | 6 | 0 |
| White | 22 | 7 | 15 |
| Years as a Postdoc |  |  |  |
| Less than 1 year | 16% | 13% | 18% |
| 1 - 2 years | 22% | 20% | 24% |
| 2 - 3 years | 25% | 20% | 29% |
| 3 - 4 years | 16% | 13% | 18% |
| 4 - 5 years | 13% | 27% | 0% |
| 5+ years | 9% | 7% | 12% |
| How likely are you to pursue a research-intensive faculty career? |  |  |  |
| Very Likely | 34% | 60% | 12% |
| Somewhat Likely | 31% | 20% | 41% |
| Neutral | 13% | 13% | 12% |
| Not Very Likely | 16% | 7% | 24% |
| Not Likely at All | 6% | 0% | 12% |

We used an ecological systems approach to examine mentoring barriers. This approach recognizes that behavior change, well-being, and outcomes are influenced across systems (with the social and cultural dimensions of an individual’s environment being the broadest system) [36–38]. These dimensions or systems can be organized by macro-, exo-, meso-, and micro-systems [39]. Thus, there are interdependent factors of both individuals and their environments that underlie behavior and outcomes, and in this case, mentoring outcomes. Our modified social ecological model posits that mentoring outcomes are first affected by an individual’s attitudes, beliefs, values, and past behaviors (**Fig 2**). Personal background characteristics such as gender, race, ethnicity, and age may also affect mentoring outcomes. Then, at the interpersonal level, mentoring outcomes are influenced by personal relationships such as those between a PhD advisor and mentee (dyadic), between two trainees (peer), or through a network such as the National Research Mentoring Network [40]. The professional relationships and social networks that a mentee takes part in have great potential to impact behaviors and outcomes. The institutional level of this social-ecological model focuses on the departmental, institutional, and organizational-specific barriers that influence mentoring practices. Finally, policies, practicies, and norms that are observed and/or perpetuated on a national or global level make up the systemic level. We examine postdoc-identified mentoring barriers on these four broad levels to better codify how structural mentoring barriers can permeate down to institutional, interpersonal, and individual levels (**Table 2**).

**Fig 2.**
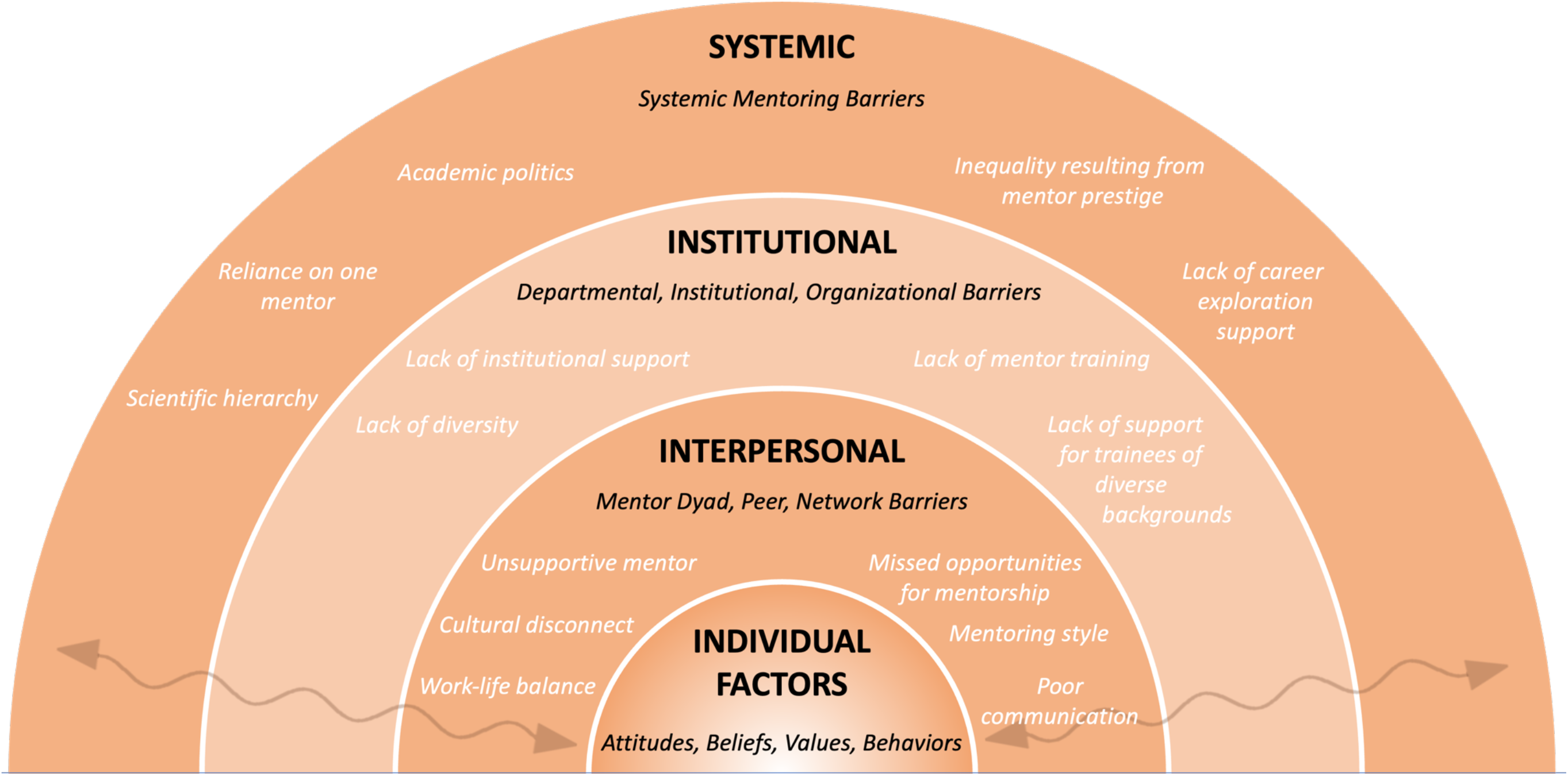
A Social Ecological Framework for Mentoring Barriers. Individual-level factors that influence mentoring outcomes are nested within mentoring relationships (dyads, peer mentoring, mentor networks), which are nested within the larger departmental, institutional, and organizational barriers. Macro-level (systemic) factors, such as lack of career exploration supports in training paradigms, may influence institutional standards and policies, which may affect the dynamics of mentoring relationships, which ultimately affect individual attitudes, beliefs, values, and behaviors (represented by the bi-directional arrows).

**Table 2.** Mentoring Barriers.

| Theme Type | Theme | Code |
| --- | --- | --- |
| Individual (attitudes, beliefs, knowledge, behaviors) | Nurturing mentors | Accustomed to nurturing mentors |
|  | Vicarious learning | Poor vicarious experience influenced career choice |
|  | Vicarious learning | Impacted by negative mentoring experiences |
|  | Vicarious learning | Success of mentor can influence confidence and beliefs about one's own potential for success |
|  | Disadvantaged backgrounds | Trained at less prestigious institutions |
|  | Disadvantaged backgrounds | Discriminatory institutional policy around maternity leave |
|  | Motivation | Desire to make mentors proud |
|  | Motivation | Persisted through the lack of strong mentors |
|  | Motivation | Fear of revealing non-academic career interests |
|  | Work-life-community balance | Familial values impacted by work-life balance |
|  | Work-life-community balance | Conflicted about proper balance between research and science/community outreach |
| Interpersonal (dyadic, peers, networks) | Missed opportunities for mentorship | Missed funding opportunities |
|  | Missed opportunities for mentorship | Not all PIs know how to be good mentors |
|  | Missed opportunities for mentorship | Mentor did not help with professional growth |
|  | Unsupportive mentor | Jealous, unsupportive mentor sabotages trainees |
|  | Unsupportive mentor | Burden of lab management responsibilities due to lack of direction from mentor in a small lab |
|  | Unsupportive mentor | Boss, not a mentor |
|  | Mentoring style | Different mentoring styles |
|  | Mentoring style | Poor co-mentorship environment |
|  | Cultural disconnect | Culturally disconnected |
|  | Cultural disconnect | Disconnect between older and younger scientists |
|  | Work-life balance | Expect postdocs to work overtime with little consideration about work-life balance |
|  | Work-life balance | Managing mentor expectations about work-life balance |

|  | Poor communication | Poor and miscommunication |
| --- | --- | --- |
| Institutional (departments, institutions, organizations) | Lack of institutional support | Lack of institutional support |
|  | Lack of mentor training | Lack of formal training for faculty roles |
|  | Lack of diversity | Lack of diversity in leadership positions is discouraging |
|  | Lack of support for trainees of diverse backgrounds | Lack of support for students from diverse backgrounds |
| Systemic (social and cultural norms, policies) | Academic politics | Managing correspondence with editors in publication review process |
|  | Academic politics | Managing correspondence with program officers in grant submission process |
|  | Inequality resulting from mentor prestige | Prestige of mentor creates unfair advantages |
|  | Lack of career exploration support | Mentors lack experience or resources to support non-academic career exploration |
|  | Reliance on one mentor | Reliance on one mentor |
|  | Scientific hierarchy | Limited academic freedom due to scientific hierarchy |

### Individual Level

#### Barrier: Access to mentors

The characteristics and experiences of the mentees involved in a mentorship relationship can shape the effectiveness of the mentoring. Thus, narrative responses from postdoctoral research participants were examined to understand how mentoring experiences were impacted by an individual’s attitudes, beliefs, values, and behaviors. Participants, particularly from the underrepresented focus groups, highlighted the challenges of being a woman, immigrant, first-generation, or underrepresented scientist. One postdoc, for example, focused on the challenges of being a woman in academia and facing institutional policies that denied access to emails while postdocs were on maternity leave, but not postdocs on paternity leave. That postdoc went on to say,

> *“I’ve kind of already alluded to [not having] really strong mentors throughout my life. I guess, for me, I’ve come this far, and I’ve done it pretty much on my own.”*

Other female postdocs reaffirmed the challenges of being a woman in science. Similarly, first-generation scientists also noted how their background did not afford them opportunities for great mentorship:

> *“I come from not a very privileged institution. When I did my undergrad, I did it in a public university in Mexico. So I’ve been raising up on that. Now, I’m here in a hospital, in a children’s hospital working. And every step of my career has been going from something less to something bigger…So, no, [mentors] haven’t handed me anything. And they haven’t actually helped me on anything. It has been me.”*

These postdocs persisted through the lack of strong mentors present in their educational and scientific journeys. However, other postdocs from racially/ethnically underrepresented backgrounds specifically sought out mentors also from URGs. These URG mentors served as role models (in addition to mentors) and fostered a strong motivation for persisting in academia. Subsequently, there is a strong desire to ‘make these mentors proud.’

> *“I definitely don’t want to let down any of the mentors that are in my life. So by keeping on the path that I’m aiming to pursue I think I can make them proud. So I want to succeed not just for me, maybe 90 percent for me but, also, to let them know hey, look, you made the right choice of betting on me.”*

We expand on this more in the mentoring solutions section below. A pattern expressed by many of these postdocs was the interest in mentors who were more nurturing. These are mentors who give positive feedback, help build a mentee’s confidence, and have patience and understanding. One postdoc from an URG noted that she joined the lab of a new URG principal investigator (PI) and was surprised when he was not as nurturing.

> *“And then as a postdoctoral fellow, I joined the lab of a new PI who is a URM and it’s actually been really challenging in the sense that he is--he will be a really great PI someday, but when I first joined he was just starting and he is very motivated and that motivation can be very grinding. He’s not somebody who gives a lot of positive feedback and he’s somebody who’s constantly reigning me in. I was so used to somebody who nurtured any time that I wanted to do something. But, in some senses, what’s been great about it is he has given--he’s really pushed me to write grants. And he’s really helped me improve as a public speaker, things that I really needed when I go speak at large conferences.”*

These comments show that the experiences from underrepresented postdocs have shaped mentoring outcomes in several ways. For some underrepresented scientists, aspects of their identity and background (e.g. being a woman or a first-generation student) affected access to strong mentors. This was not expressed among the postdocs from WRGs. Unlike their well-represented counterparts, postdocs from URGs struggled to find mentors willing to take the time to provide strong mentorship. They had to then persist through the lack of strong mentors in order to reach higher academic stages. For other scientists from URGs, they were attracted to and sought out more URG mentors, who were often nurturing in their mentorship style.

#### Barrier: Vicarious experiences that lead to poor mentoring outcomes

Vicarious experiences allow individuals to learn from the experience of others. One postdoc noted how watching her mentor struggle with attaining grants, in-part influenced her future career choice:

> *“So I came in two years ago starting my postdoc with the intention of doing everything I could to get into an academic position. And between not getting any of the six grants that I applied to and watching my advisor struggle through trying to apply for tenure despite knowing that she’s a phenomenal mentor and a really great scientist and she was awarded it but just the stress of that and the amount of work and everything I just kind of decided that that might not be career path for me.”*

Some mentors may feel helpless at the thought of mentees interpreting their struggles negatively. Thus, it is important to take an active role in managing the expectations of what it means to be a researcher in academia for trainees. Another postdoc noted how the success of her mentor influenced her confidence and beliefs about her own potential for success.

> *“I had a lot of trepidation about pursuing a research focused faculty appointment versus a teaching focused faculty appointment at my former institution because my mentor was struggling to get RO1s funded, and it was just scary all the time. And now I’m in a postdoc position with someone who’s very well-funded and has really mentored me in grantsmanship, and I feel a lot more confident about pursuing that sort of grant funded research career because I see that it’s possible.”*

Other postdocs have noted how negative mentoring experiences overall impact their own approach toward mentoring:

> *“So my mentoring experience up until my postdoc really hasn’t been very positive; if anything it’s kind of hurt me, but at the same time it’s also helped because it was to the point that other faculty at the university just knew how bad it was What I took from that is how not to be a mentor. So just everything, what not to do, and how not to run your lab.”*

These quotes highlight the impact that previous vicarious and/or negative mentoring experiences have on both future mentoring relationships and career outcomes. They also reinforce the influence that one’s background, beliefs, and prior experiences have on the mentoring relationship and future mentoring practices.

### Interpersonal Level

#### Barrier: Research advisors who lack good mentoring practices

Mentoring barriers at the interpersonal level were largely characterized by personal relationships between a research advisor and mentee (dyadic). In one case, a postdoc characterized a research advisor as jealous, unsupportive, and actively preventing growth and advancement. However, in most of the comments, the focus group participants noted that mentors were not actively creating barriers to their success. Rather, there was simply a lack of good mentoring or missed opportunities for mentorship. One postdoc highlights:

> *“You know, I’m sort of kicking myself after I left, because there were clearly all of these opportunities for, like, a URM supplement for his R01, and all these other things that I don’t even know if he--I mean, I’m sure he didn’t take advantage of it, but there were all of these missed opportunities that he just was not interested in even pursuing, because he was just not interested in having me write anything.”*

There was an agreement that some PI’s do not know how to be good mentors. They did not help with professional growth or create opportunities for advancement. One postdoc described his PI as a “boss, not a mentor.” Others noted a misuse of their position and time by mentors, such as a burden of lab management responsibilities:

> *“…It’s just myself, the one postdoc, and one research technician. So, a lot of responsibility falls on me in terms of that, running the lab, and also with grants, and all that. So, it just extends to a lot of--a little bit irresponsible and unorganized mentorship where it’s not very helpful and healthy for me.”*

Underneath the missed opportunities for good mentoring practices were specific issues such as misaligned expectations or poor communication.

> *“So, I think that right now in my position, my current mentor is not supportive and does not provide any mentorship or even a lot of scientific feedback. But his mentorship is also a little bit different because he thinks he’s helping and doing right in certain things. And the way that he communicates, it comes out very negative, but I think because of the passive-aggressive behavior in terms of, like, saying that if I’m not going to publish, then I might not get re-hired and then giving examples of things that are not necessarily very responsible and ethical. So, that’s one of the concerns that I’ve had recently. And then trying to see if I can actually mend and work on that relationship, but I’m not getting much back from it.”*

In some cases, there were misaligned expectations around work-life balance that created challenges in the mentoring relationship. The postdocs found certain expectations from their mentors very discouraging if those expectations included longer-than-normal work hours and conflicts about time spent in the laboratory. For example, one postdoc noted:

> *“I am constantly reminded that I should be working from seven in the morning till two in the morning, which was what they did when they were postdocs. And, realistically, as a female, I don’t feel safe wanting to leave lab every day after two in the morning when I would take public transportation.”*

Other misalignments were over opposing mentoring styles and fit:

> *“I’ll say I think I’ve had really excellent mentors both now in my postdoc and during graduate school. My current postdoc advisor is well-established. She’s trained many students and postdocs who have gone on to have academic careers. And so she clearly has the ability and the name to be able to do that. Although, she--and I don’t think that she should do this, but she’s not necessarily going to like push me and say like, ‘Hey, you need to do this. You need to do that,’ because, you know, it has to be like an intrinsically motivated thing especially for this [academic] career path.*

> *I would say in contrast, I worked with a more junior faculty for my graduate career and she was a little bit more of the mindset that if you didn’t stay in the academic environment that you were sort of failing. She wanted us all to kind of grow up and be her. So even if she didn’t necessarily have like that name or all the tools to be able to like usher us into an academic career, I think she that idea for all of us.”*

Similarly, another postdoc noted contrasting mentoring styles in their mentoring experiences:

> *“Well, my first mentor in grad school, he was very hippie. So he had a very nice attitude. I learned a lot about how to approach other people and how to deal with other people. But he was not very good on finding new opportunities. He was in a very secure position, so he was not very adventurous. So when I moved to my second position my advisor here, he’s a dreamer. He just dreams but everything he reads, making a new project, and making a new idea. And with him we need to kind of find a feasible idea that we can actually pursue. So they have their own personalities and I have my personality…I don’t think I am either the hippie, my first advisor was, or the type of dreamer that my new advisor is. It’s just you’re on your own.”*

#### Barrier: Culturally disconnected mentors

The lack of cultural awareness or cultural humility was a common theme among the underrepresented focus groups. Some mentors, although well-meaning, do not know how to approach cultural differences and can come off as not being supportive. Other times, they can be insensitive and come off as offensive. One postdoc describes comments from a mentor that were received as derogatory.

> *“Thankfully the mentor I have now grew up on a farm in the middle of nowhere Missouri, so he pulled himself up by his bootstraps. But a previous co-mentor like his father was a faculty member at Berkeley and then his grandfather was a faculty. So that was a silver-spoon-in-mouth situation. So there were always snide comments here and there that you can interpret it as being derogatory. So it depends on the person. But I think, on the whole most people, just don’t care, just publish and get grants, move on.”*

Another interesting point that arises from this particular quote is that this postdoc found commonality and shared values in the background of her current mentor who helped himself succeed without the privilege of wealth and resources, as noted in contrast to the previous mentor. Disconnect is not limited to just cultural backgrounds. Another focus group participant highlighted the disconnect between older and younger scientists:

> *“I don’t feel like necessarily they’re thinking about what we should be--how each path’s a little bit different and independent from what it was for them and their training maybe, like, 20 years ago. And, so, there’s a disconnect I still feel that with the younger scientists coming in and then us still finding our path in academia. I feel that that’s also something that I’m thinking that other people that I’ve talked to have experienced that.”*

Both examples point to a significant barrier in cultural awareness and engagement that can exist for many postdoctoral fellows, especially from URGs.

### Institutional Level

#### Barrier: Lack of institutional support for postdocs

According to the focus group participants, many institutions fail to provide services, training programs, support, and attention to postdoctoral fellows. In addition, postdocs point out the need for formal training for faculty roles:

> *“I actually had a pretty decent graduate career, published pretty well, publishing pretty well in my postdoc and I am terrified out of my mind to even think about the faculty position. We really don’t have any sort of training before a faculty position, you’re told to publish while you’re a postdoc, maybe get some grants, interview for faculty and then you’re thrown to the sharks to really fend for yourselves. And if you’ve had some good mentors, they can provide guidance, but I mean at that point you’re pretty much on your own, running your own business, you’re accountable for people now, you’re accountable for their future, their education as well as your own.”*

They highlight the need for more institutional support to transition into faculty careers, which has arisen in previous studies [5]. A lack of mentoring was also noted as a key part of the lack of institutional support.

#### Barrier: Lack of support for trainees from diverse backgrounds

Multiple participants of one particular focus group highlighted the lack of diversity in leadership positions and the lack of support for trainees from diverse backgrounds (underrepresented groups):

> *“But I also see it as a place where much of the leadership are professors who have been there for a very long time in their careers and, so, a lot of them are faculty, white men, in those positions making a lot of decisions [about] the way that they’re running departments or their certain field of research. I find that discouraging, the lack of diversity and support of diverse students at the PhD level or at the postdoc level.”*

In their opinion, the lack of diversity in leadership positions has led to decision making and views on career outcomes that have not evolved with the views of trainees. According to the postdocs, these types of barriers at the institutional level have led to a culture characterized by a lack of support and mentorship. Moreover, this culture is upheld by the leadership when little to no intervening occurs. It is similar to the mentoring gap previously described in the literature [41, 42]. For marginalized trainees, access to mentoring networks is often more limited than that of their peers. Not everyone has equal access to mentors. This mentoring gap is frequently associated with a person’s background, their race, class, and gender, and perpetuated by structural inequities. We expand on this phenomenon, pointing out the structural barriers that maintain the mentoring gap.

### Systemic Level

#### Barrier: Academic politics

At the systemic level, postdocs focused on mentoring barriers such as ‘academic politics’, scientific hierarchy, and the structural reliance on one PI for career success as a postdoc. One postdoc noted,

> *“Well, in my experience, I feel I have the skills to do research, to design experiments, to teach graduate schools how to perform research. Still what I find difficult is this: managing the politics of, for instance, when you get a paper reviewed and then you have to reply and then how to deal with reviewers and with editors. And the same with funding how when you get your grant application gets reviewed, the correspondence with the program directors and the staff that make the decisions to fund your research or not. I find myself that I need mentoring in that aspect.”*

Managing correspondence with editors in the publication review process and program officers in grant submission process is not something that is formally or systematically taught to postdoctoral fellows. One must usually rely on the mentorship of the research advisor/PI to teach these skills if not already learned. The postdocs felt that this structure limited their academic advancement. Scientific training hierarchy and power dynamics were also cited as a barrier, where a younger trainee’s ideas were stifled.

#### Barrier: Reliance on one mentor

An ‘over-reliance’ on one mentor was problematic for the postdoc participants. One postdoc noted,

> *“I think there’s one thing regarding mentorship that I’d like to add. One of the things in grad school that’s kind of tough is that unlike in a corporation where you have sort of coverage from your bosses, when you’re a PhD student, you’re at the whim of your PhD mentor and you don’t have a lot of security and support beyond that so if you have problems then it’s not like you have other people who can support you in furthering your career, they’re the one person you have to rely on and that can be a really hard thing.”*

The impact of mentor (and institutional) prestige was a common theme across focus groups. If mentors held a certain prestige in the scientific community, they could often create seemingly unfair advantages for certain trainees. One advisor, just based on their name, could have large influences on a trainee’s career trajectory. For example, this postdoc notes:

> *“…he’s well known in the Alzheimer’s field so the way he helped me is not even active, it’s just passive. It’s just from his name. The fact that I postdoc under my mentor, like what you were saying, [other participant], about the university. Well, also, the mentor is like the same, “Oh, you came out of So-and-So’s lab. Well, then okay, you can get whatever position wherever,” which they don’t have to do anything for. I mean they’ll write a letter of recommendation but it’s just the name, really.”*

#### Barrier: Lack of support for career exploration

Finally, the postdocs stressed the importance of mentor training and resources. Postdocs feel unsupported by both their institutions and their mentors as they explore their career options after their postdoctoral training. If they don’t find resources through their institution, they often turn to their mentors. Yet, many mentors lack the experience or resources to support non-academic career exploration. For example, one postdoc said,

> *“…I think mentors in academia they are too focused on academia. This is what they have been doing for all their careers, so they don’t contemplate other career pathways. Or if you are trying to get into something they probably won’t help you at all because they don’t have that experience and they don’t contemplate the other options, in general.”*

### Mentoring Solutions

In addition to codifying the barriers that postdoctoral trainees face in their mentoring relationships, we present mentoring solutions to overcome many of those barriers. These proposed solutions were developed by the postdoctoral participants and organized into four groups: (1) mentoring solutions in diversity, equity, and inclusion, (2) encouraging and helping trainees to develop mentoring networks, (3) promoting motivation, and (4) building skills and support (**Fig 3** and **Table 3**). The format used in Fig 3 can serve as a printable toolkit or reminder for mentors as they interact with all of their trainees. Below, we highlight and describe in detail these mentoring solutions.

**Fig 3.**
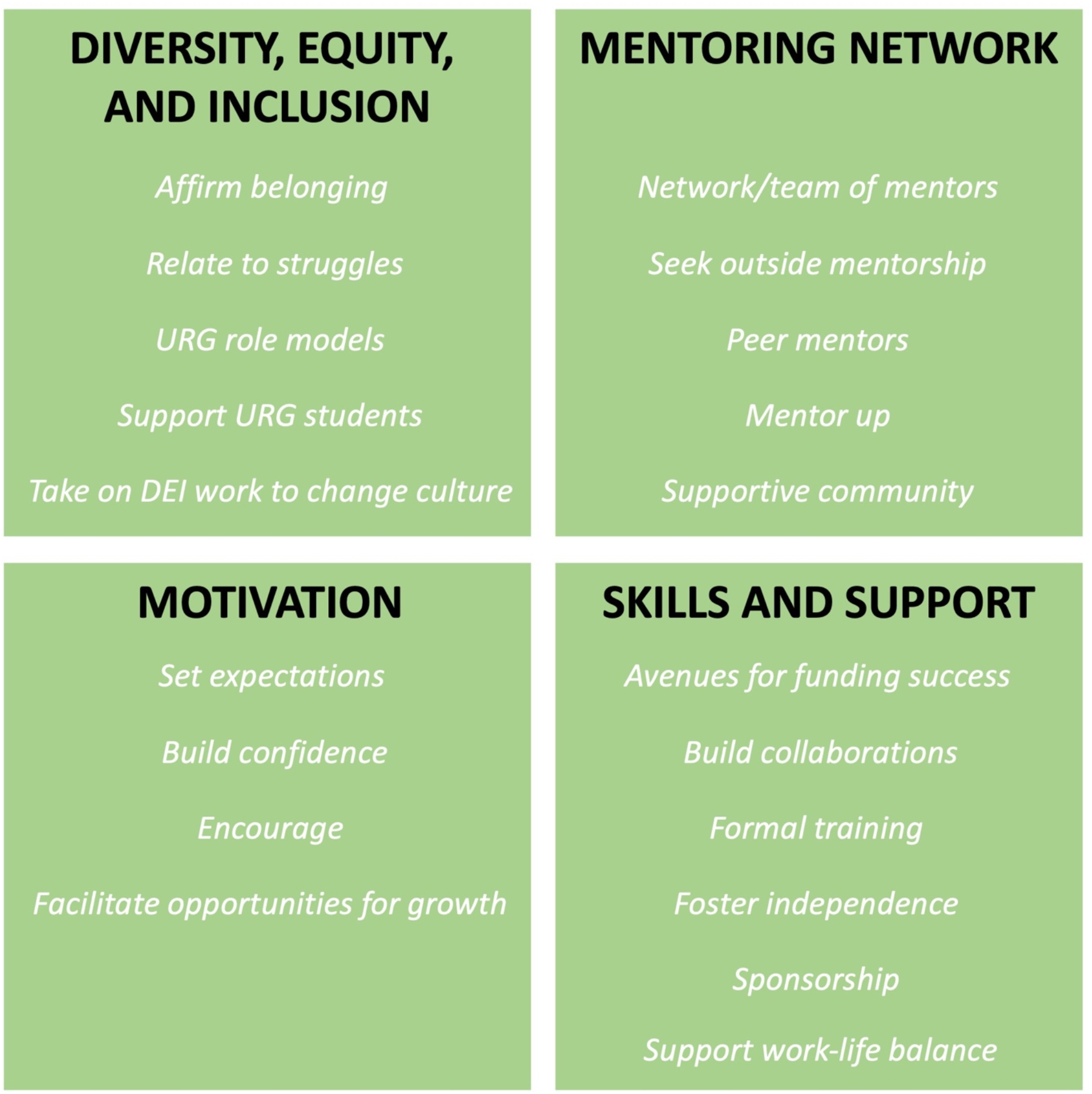
Mentoring Solutions Proposed by Postdocs. Postdocs proposed solutions for overcoming mentoring barriers, organized into four themes: (1) mentoring solutions in diversity, equity, and inclusion, (2) encouraging and helping trainees to develop mentoring networks, (3) promoting motivation, and (4) building skills and support.

**Table 3.** Mentoring Solutions.

| **<u>Mentoring Solutions</u>** | **<u>Theme</u>** | **<u>Code</u>** |
| --- | --- | --- |
| Diversity, Equity, and Inclusion | Affirm belonging | Mentor affirms URG student's belonging |
| Diversity, Equity, and Inclusion | Relate to struggles | URG mentor relates to mentee struggles |
| Diversity, Equity, and Inclusion | Support URG students | Support URG students with mentorship, professional development, and belonging |
| Diversity, Equity, and Inclusion | Support URG students | URM mentor not nurturing but helped development in other ways |
| Diversity, Equity, and Inclusion | Take on DEI work to change culture | Participate in diversity and inclusion work to change academic culture |
| Diversity, Equity, and Inclusion | URG role models | Increase diversity in faculty and leadership |
| Diversity, Equity, and Inclusion | URG role models | URG mentor motivates desire to continue in science and increase diversity |
| Diversity, Equity, and Inclusion | URG role models | Motivated by first generation and mentors of color outside of academia |
| Mentoring Network | Network/team of mentors | Develop a team of mentors |
| Mentoring Network | Network/team of mentors | Create a network of mentors |
| Mentoring Network | Network/team of mentors | Develop diverse mentorship network early in career |
| Mentoring Network | Network/team of mentors | Have honest conversations with professionals across diverse fields early |
| Mentoring Network | Network/team of mentors | Unofficial mentors and the importance of a community of mentors |
| Mentoring Network | Seek outside mentorship | Sought outside mentorship to supplement in other areas |
| Mentoring Network | Seek outside mentorship | Go outside of traditional channels for mentorship |
| Mentoring Network | Seek outside mentorship | Honest advice from diverse network to help with career development |
| Mentoring Network | Peer mentors | Peer mentors outside of discipline |
| Mentoring Network | Peer mentors | Peer and near-peer mentorship |
| Mentoring Network | Peer mentors | Peer mentorship |
| Mentoring Network | Mentor up | Develop skills necessary to get the most out of mentors |
| Mentoring Network | Supportive community | Look for an established, supportive community |
| Motivation | Expectations | Help mentee exceed expectations of themselves |
| Motivation | Build confidence | Build up mentee's confidence |
| Motivation | Build confidence | A mentor who builds up your confidence helps fight imposter syndrome |
| Motivation | Encourage | Encouraged by mentor |
| Motivation | Prioritize mentorship in lab choice | Prioritized mentoring in choosing a lab |
| Skills and Support | Avenues for funding success | Mentor provides avenues for funding success |
| Skills and Support | Build collaborations | Establishing new collaborations with other investigators |
| Skills and Support | Build collaborations | Facilitate opportunities for growth and networking |
| Skills and Support | Formal training | Better training for faculty positions |
| Skills and Support | Foster independence | Help mentees gain independence through varied training and mentorship |
| Skills and Support | Sponsorship | Mentee benefits from mentors' reputation |
| Skills and Support | Support through processes | Mentorship during faculty and interview process |
| Skills and Support | Support to pursue other interests | Mentors supported mentee in pursuing other interests |
| Skills and Support | Support work-life balance | Support for time off, childcare, tenure clock |

#### Solution: Affirm belonging

Trainees from URGs progress through academic pipelines often with feelings of not being fully welcomed or valued at their institution [43, 44]. This can reinforce negative stereotypes and reduce self-efficacy to the point where URG trainees can begin to question their potential or motivation [45]. Our URG focus group participants highlight the need for mentors to affirm trainees’ sense of belonging:

> *“So, the perfect example, I think, was--I don’t know if you all remember when Justice Scalia was talking about how black students shouldn’t go to top-tier schools for science, that they should stick with lower-tiered schools. And this was all over the news and she called me into her office, and I was like, ‘Oh, God, I’m in trouble.’ And she says, you know, ‘I just want to let you know I heard everything in the news, but I just wanted to let you know specifically you belong here.’ And--I don’t know--that was just a turning moment for me, where sometimes we hold things, especially as you’re around students that maybe our peers don’t hold in terms of just baggage [or] micro-aggressions or whatever it is. And so, just hearing that support that was outside of the realm of research was just really great for me.”*

Similarly, postdocs highlight the need to support URG students with mentorship, professional development, and belonging:

> *“There’s just a lack of diversity up that pipeline. And the further of you get the less diversity you see. And, so, for me, you know, I absolutely agree that one way to change it is to be in the research institutions and be that face that other--that students can see and aspire to, but the other aspect of it is the administrative side. And I think that really lending support, both financially, both mentoring, professional development, just having that sense of welcoming to a lot of students to traditionally under-represented groups is incredibly important in their graduate careers.”*

#### Solution: Participate in diversity and inclusion efforts

The previous quote also highlights the need for mentors to take on and participate in diversity and inclusion efforts themselves. For example, joining committees, recruiting URG trainees, and conducting outreach efforts are important and can be impactful, but postdocs stress the need for mentors to think about building a more inclusive academic culture, which is often overlooked.

> *“There’s a lot of things that are not inclusive in academia. There’s a lot of things that diversity affects that, I think, need to be changed. And, so, I’m actually a part of a training and inclusion workforce here that I help run and there’s a lot of things that people are unhappy with and a lot of things why the retention is so low and a lot of reasons why people get their PhDs, but then you don’t see them in postdoc, a lot of reasons why they go somewhere else. And a lot of it is just because you don’t feel welcomed. Right? You don’t feel the environment you need. I mean there’s a lot of things I think that need to be changed, like [Participant 1] said. And if I stay in research, if I stay in R01, of course, I love what I do. I love my science. I have the independence that I need, but I’m hoping to also make a difference, too, and just the entire environment, because I don’t know who else will I guess.”*

#### Solution: Hire more URG role models

Another solution that was emphasized was the impact of URG mentors on URG trainees. URG mentors can relate to and understand the struggles of URG trainees. These are qualities that can be found in and practiced by non-URG mentors, as well. Postdocs note that URG mentors also motivate URG trainees to continue in science, thereby increasing diversity.

> *“So, for me, I wanted to do a postdoc because I had a fantastic graduate advisor. He was--he’s Latino and he was very much if you’re a minority, you should stay in science, because there’s not a lot of us. So, my goal is ultimately to be a PI, so I can try to pull more under-represented minorities up into scientific fields, because there really aren’t that many.”*

This demonstrates that there is a need to increase the number of URG faculty, but trainees are also motivated by URG mentors outside of academia, particularly those in leadership positions.

> *“I think what I’ve been--I feel really highly motivated is by the mentors that I have made outside of science and research necessarily, having the opportunity to actually network with a lot of different people and scientists and different-- and social scientists and actual traditional biomedical scientists through different organizations and those opportunities. And I feel really highly motivated by seeing other first generation and People of Color taking on these leadership positions.”*

#### Solution: Create a network/team of mentors

One of the most vocal recommendations that the participants offered was to create a team or network of mentors.

> *“And to reiterate on the mentorship theme that [the other participant] was mentioning, find yourself a mentorship team. I think that’s really important to have many people that you can approach, and it doesn’t have to be the same person--you don’t have to go to the same person for grant-writing support that you do for work-life balance support. I think that it’s important to have a whole team of mentors at your disposal.”*

There was a lot of encouragement to develop a diverse mentorship network early in one’s career and have honest conversations with professionals across diverse fields. Honest advice from a diverse network of mentors will also help with career development. One postdoc also highlighted the importance of a community of unofficial, or “mini” mentors who are often your peers:

> *“I think the term ‘tribe’ was used at some point, your community, that it’s--I think of the adage that it takes a village to raise a child. And sometimes it’s leaning on those sort of unofficial mentors or mini mentors. Sometimes it’s a peer that takes on very briefly a mentorship-type role for a moment, for a conversation. And that sounds like it’s sort of reaching and being poetic, but it’s really not, especially in the absence of, I would say, comprehensive, competent, official mentorship. It’s just that’s how you receive mentorship.”*

Similarly, another postdoc stressed the importance of an established, supportive community:

> *“I actually--totally agree with you, actually, because I think that such a huge part of my success has been the support of an entire community. I mean, genuinely people have just come out of the woodwork to help me manifest all of my dreams and I mean that truthfully. And, so, I guess, for me right now, what I do when I go visit universities is I am looking for a community like that, that I don’t have to necessarily build. I’m looking for communities that are already there and community-based science and collaborative science and people who see me holistically, not just as a researcher, but as somebody who is also passionate about teaching and mentoring and also somebody who’s very passionate about outreach and I’ve just been--yeah. I’ve been really lucky and I have confidence that with--in the right environment I’ll be successful.”*

#### Solution: Seek mentorship outside of primary mentor

Like building a community or network of mentors, the postdocs encouraged seeking mentorship outside of one’s primary mentor or even their department.

> *“But the challenging-- and I guess this is where a greater sense of community comes in, people from my department, from other departments at the university, just because I did my postdoc at the same university I did my PhD at--people have stepped up to mentor me and provide guidance and help get me opportunities, and really continue to allow me to take a seat at the national stage as well. And, so, I guess the lack of mentorship I’ve seen in my postdoc, I’ve been able to supplement in other areas from other people.”*

Seeking external mentors may involve some self-reflection regarding one’s gaps in skills or training. The postdocs encouraged going outside of traditional channels to get the appropriate mentorship or guidance, when needed.

#### Solution: Seek out peer mentors

Another emphasized recommendation from the focus group participants was the value of peer or near-peer mentors:

> *“I actually get a lot of mentorship from colleagues who are postdocs who are a little bit--in assistant professorship positions right now and that’s where I’ve been meaning to look at. Even though they’re not in the same discipline I’m in, I’ve actually been learning a lot from them.”*

Others agreed with the notion to seek out peer-mentors, especially for career development:

> *“Yes, I think in my case I could name a few good colleagues that were, more or less, at my same stage that I was, and they were also sharing with me his or her perspectives on how they were doing and what was the plan for later. So, I think that they really act as mentors.*

> *Although as [the other participant] said, like, not officially maybe, but they also--you know, try to understand and try to help with personal advice.”*

#### Solution: Mentor “up”

Postdocs also advised to develop the skills necessary to get the most out of mentors.

> *“I think that as trainees there are skills that we need to develop in order to get the most out of our mentors because some of them are not naturally inclined that way.”*

#### Solution: Build up mentee’s confidence

It has been shown previously that higher research self-efficacy is associated with larger numbers of publications and a greater intention to pursue research-intensive careers [14]. Postdocs have also highlighted the importance of building up a mentee’s confidence and helping mentees exceed even their own expectations of themselves. One postdoc notes:

> *“I think that’s probably the most important thing a mentor can do is to build up your confidence and let you know, “Hey, you are smart enough to do this. You can accomplish such and such.” And towards that respect I had the opportunity of working with a Nobel Laureate when I was an undergrad, and he started that process for me. We published and he offered me a position in his lab, although, it was at [University]. I couldn’t pick up and go. But, you know, him having the confidence to say, “Hey, I know you’re not accepted here but if you want I would like to have you as a graduate student here,” really kicked things off. And then my graduate mentor he’s also saying, “Hey, you’re a really smart man. You can do this and that.” So even though it’s not--I don’t know if that’s more than what they’re expected to do but the confidence building definitely is a good thing if you can have that.”*

This postdoc went on to note the importance of confidence building in fighting imposter syndrome.

> *“[A mentor who builds up your confidence] is definitely one of those things where if you ever have that imposter syndrome, you’re like, well, these really, really smart people believe that I’m good enough to do it. And my track record says I’m good enough. So it sort of helps you get through those moments of self-doubt and that’s very important.”*

Another postdoc described where their mentor encouraged her mentees to think freely and develop their own ideas.

> *“I had a really similar experience during grad school where my mentor was absolutely fantastic, she treated everyone as a postdoc, everyone has their own opinions and was encouraged to think on their own.”*

#### Solution: Help provide avenues for funding success

Grants and funding are crucial for research success in academia. Postdocs note that the mentors who support their mentees in applying for funding and encourage learning how to successfully acquire grants were the most helpful.

> *“My advisor is really helpful. I think it’s what she knows, you know, as an academic, that’s what kind of happens. But she was very helpful in helping me find grants and apply for grants and writing papers, you know, kind of not even really pushing me necessarily but kind of giving me avenues and directions of which way to go towards academia.”*

Similarly, establishing new collaborations is important for long-term success. It was noted by one postdoc that establishing new collaborations was something that you have to “learn by doing it.” Others noted that their mentors aided in this process, and this was valued:

> *“If my PhD advisor had an old data set that he hadn’t done anything with, he would say, ‘Hey, you want to write this up?’ He would find opportunities for me to meet new people and collaborate with people.”*

#### Solution: Better training and preparation for faculty positions

There seems to be a dire need for better training and preparation for faculty positions, as well as the application and interview process. Whether this is in the form of more formal training opportunities or workshop series, or managed through individual mentors, postdocs highlighted the impact that this could have on their success:

> *“I actually think that it’s almost impossible to go down a faculty interview or apply for faculty position without having great mentors, having that experience behind your belt and having them really guide you through the process because it is so daunting and so exhausting that really without their help, it’s nearly really impossible.”*

#### Solution: Support mentees in their career development

One particular postdoc described her exploration and interest in the field of public health, although her PhD training was in the basic biomedical sciences. She talked about the support that her mentors offered her in exploring this route, which led to a unique faculty position that will allow her to combine both fields in a very translational research setting.

> *“Both as a PhD candidate and now, I’ve had really supportive mentors that have let me from the beginning ask my own questions and, as a PhD student, explore my interests that were not purely basic science. And, even then, I was able to start dipping my toes into public health and then, similarly, always letting me have that opportunity to do the outreach work and to get into the--start getting into the communities. And, even now, despite not--I guess giving me the time to develop relationships in the community here as a non-[western state] or [western city] native was super important for the type of work that I was--that I’m interested in doing moving forward, and to give that time and understand that I--you know, that if I am going to be successful that I need to know how to manage my time and do all the things and having confidence that I could do it has been the most important thing and just the--yeah, their support is the number one thing.”*

#### Solution: Offer support for time off, childcare, tenure clock

Another unique postdoctoral experience that highlights structural challenges, especially for women, is described below. This postdoc, who had just accepted an offer for a faculty position, started her PhD program later in life. She describes how this was actually a more suitable time for her to develop her career due to the general lack of support that academia offers in terms of time off, childcare, and tenure clock. This resonated with other members of the focus group.

> *“I actually have avoided a lot of these and I think the reason is because I went to school at 41 and I think a lot of these problems have to do with the very, very rigid timeline of PhD, postdoc and faculty and that clock is ticking during your prime earning years and your prime reproductive years especially for women and I think it’s just a collision course, there’s just no way you can do it all simultaneously and it’s very stressful. And I think that having had a career in my 20s where I was able to build up equity and a 401[k) and buy real estate because I was making an actual salary and then once I was settled to go back to school took all the pressure and I had my children already. So I think that these are all things that if there were more flexibility built into the system time wise, better leave, more childcare support and that tenure clock which is just, it’s just so rigid, it would help a lot especially for women.”*

## Discussion

This study examined the experiences of URG and WRG postdoctoral researchers and discovered the following major findings: (1) the lack of institutional diversity and mentor training are impacting mentoring relationships. (2) Broad systemic mentoring barriers can be mitigated through policies, networks, and training. (3) Postdocs, especially from URGs, seek more nurturing mentors who understand their values and can build their confidence.

It has become clear from this study that the lack of formal institutional mentor training is a structural barrier impacting mentoring relationships and outcomes of postdoctoral fellows, especially from URGs. In fact, both WRG and URG postdocs stressed the importance of this. Due to training grant requirements and *train the trainer* resources such as those offered by the Center for the Improvement of Mentored Experience in Research (CIMER), many institutions are now offering some form of mentor training. These trainings can be effective, as shown by Pfund et al. and others [46, 47]. However, large effects of mentor trainings were not reported by the postdocs in our focus groups. We think this is largely due to the medium and frequency of mentor trainings, the trainings having minimal effects on changing the culture of mentorship at institutions, and the inability of the trainings to affect the behavior of PIs with bad mentoring practices. Mentor training must be coupled with policy change, ‘checks and balances’ to protect trainees, and incentives and awards to promote good mentoring and culture change.

Similarly, a lack of diversity, particularly among faculty, continues to impact mentoring outcomes. Institutions have long been aware of this recruitment challenge, but less conscious of the exclusive environment that some URG postdocs experience. Postdocs want to be supported and treated at least as equally as graduate students. However, for URG trainees, there can be a poor sense of belonging at academic institutions due to factors such as implicit bias, microaggressions, and an overall inequitable lack of support in their training. Moreover, higher levels of mentor-mentee psychological similarity lead to higher levels of psychosocial support, relationship satisfaction, and publications, (i.e. Hispanic trainees with a Hispanic faculty mentor report engaging in more coauthoring opportunities than peers with non-Hispanic mentors) [16]. Some institutions have required diversity and/or implicit bias trainings for their staff, trainees, and faculty. Others have also created cohort programs specifically for postdocs such as IRACDA and the Carolina Postdoctoral Program for Faculty Diversity to help with both a sense of belonging and also skill development towards faculty careers. These efforts should be widely embraced among a myriad of tools to improve mentoring and diversity for postdoctoral trainees.

Our study also brought to light the importance of policies, networks, and training in mitigating broad systemic mentoring barriers. Mentoring barriers on the systemic level consistently impact barriers on the individual, interpersonal, and institutional levels. For example, there has been an increase of postdoctoral trainees pursuing nonacademic careers. For some postdocs, like the one described earlier in this study who watched her advisor struggle through applying for tenure despite knowing that she is a phenomenal mentor and great scientist, the shift to a nonacademic career is largely based on negative vicarious mentoring experiences. Upstream of this are the national and institutional pressures for teaching more courses and increasing grant funding, ultimately affected by federal funding and institutional business models. Academic structures like this systematically affect the dyadic relationship between mentor and mentee. Moreover, these structural mentoring barriers make it difficult to implement effective dyadic interventions. However, the postdocs in our study point out that mentoring outcomes from these barriers can be mitigated. In addition to policy changes, building a network of mentors will provide postdocs with more than just one primary resource for career and professional development [48, 49]. As noted by others, training for both postdocs and mentors (in the context of mentor training) will help with positive outcomes [50]. Subramanian et al. also suggest increasing collaborations between research mentors and career development educators [51].

One unexpected and striking outcome of this study was the acknowledgement by many postdocs, especially those from URGs, that they sought or appreciated mentors who were more nurturing. These were mentors who often gave positive feedback, helped build their confidence, understood their values, and had patience and understanding. For URGs postdocs, they often found these characteristics in URG faculty mentors (or other peers). Thus, there was an affinity for women or underrepresented racial and ethnic group (African American/Black, American Indian and Alaska Native, Hispanic/Latinx, Native Hawaiian and other Pacific Islander) mentors. Many URG postdocs were accustomed to nurturing URG mentors, due to the role that these type of mentors had played in their academic and scientific development. One postdoc was even surprised after joining the lab of a URG faculty member and realizing he was not as “nurturing” as she had hoped. She noted how he was not someone who gave a lot of positive feedback; however, she admitted that he kept her focused on the skills she needed to develop to be successful, such as focusing on grant writing and not getting too distracted by non-relevant (scientific or non-scientific) activities. Overall, postdocs derived a lot of motivation from nurturing mentors, and often desired to make them proud.

Not all URG postdocs sought nurturing mentors. Some noted a historic lack of access to strong mentors in their journeys and scientific development, and largely were able to ‘pull themselves up by their bootstraps’. They did not attribute their success to effective mentorship, rather to hard work alone. The common themes between postdocs seeking nurturing mentors and those who attribute their success (thus far) to hard work were the barriers and hardships that both groups have had to overcome to reach this point in their training. These hardships were not unique to URGs and were found among international postdocs, first generation, and other often marginalized groups. To address this, postdocs suggested less of a reliance on one mentor and instead encouraged building a mentor network or team of mentors to foster success. There is a growing body of literature that shows that having a personal network of mentors improves retention in research careers [4, 52]. What is not clear, however, are the aspects of a mentoring network that are critical for the success of underrepresented early career researchers, and how URGs break into or create these networks.

The present study has some limitations. Though participants were diverse in terms of gender, race/ethnicity, and training stage, our sample size was relatively small compared to the available pool of biomedical postdocs and did not represent the entire range of gender and racial/ethnic diversity in the US. We also were not successful in recruiting postdocs from Historically Black Colleges and Universities (HBCUs) for this study. It has been shown that investigators at minority-serving institutions often have less access to collaborators and departmental colleagues with federal funding [53]. Thus, we did not gain further insight as to the structural mentoring barriers faced at minority serving institutions. Furthermore, trainees were asked about the totality of their academic experiences rather than focusing specifically on aspects of mentorship only. Although we coded all of the data and observed data saturation, it is possible that we could have missed opportunities for postdocs to delve further into their mentoring experiences during the focus group. This work was a follow up study conducted to inform unanswered questions for a larger survey study that assessed the factors that influenced postdocs’ career decisions by race/ethnicity and gender.

In this manuscript, we discussed the major systemic, institutional, interpersonal, and individual mentoring barriers faced by postdoctoral trainees and possible solutions. Overall, postdocs point out missed opportunities for mentorship and an overall lack of support for trainees at their stage. Not all PIs know how to be great mentors, but an understanding of the larger factors affecting mentoring outcomes, as outlined here, can help to enhance the culture of effective mentorship not just for postdocs, but other trainees as well. Many of the barriers outlined in this paper are generalizable to undergraduate, graduate, and medical students. Moreover, academic institutions are not the only types of institutions that have the power to influence mentoring outcomes. We hope this social ecological approach to understanding mentorship barriers provides a framework for improved mentoring outcomes for all types of institutions.

## Acknowledgments

The authors wish to thank Dustin Thoman for the helpful comments. Research reported in this publication was supported by the National Center for Advancing Translational Sciences of the National Institutes of Health under award number UL1TR002384, and the National Institute of Allergy and Infectious Diseases (NIAID) under the award number K24 AI10732. The content is solely the responsibility of the authors and does not necessarily represent the official views of the National Institutes of Health.

